# A Guide for Quantifying and Optimizing Measurement Reliability for the Study of Individual Differences

**DOI:** 10.1101/2022.01.27.478100

**Authors:** Ting Xu, Jae Wook Cho, Gregory Kiar, Eric W. Bridgeford, Joshua T. Vogelstein, Michael P. Milham

## Abstract

Characterizing individual variations is central to interpreting individual differences in neuroscience and clinical studies. While the field has examined multifaceted individual differences in brain functional organization, it is only in recent years that neuroimaging researchers have begun to place a priority on its quantification and optimization. Here, we highlight a potential analytic pitfall that can lead to contaminated estimates of inter-individual differences. We define a two-dimensional individual variation field map to decipher sources of individual variation and their relation to fingerprinting and measures of reliability. We illustrate theoretical gradient flow that represents the most effective direction for optimization when measuring individual differences. We propose to use this general framework for dissecting within- and between-individual variation and provide a supporting online tool for the purposes of guiding optimization efforts in biomarker discovery.

## Introduction

Over the past decade, neuroscience has witnessed a shift in focus from group-level brain investigations of the average effect across individuals, to the delineation of variations among individuals and their relations to genetic and phenotypic variables (e.g., demographic, behavioral, cognitive, psychiatric)^1–7^. Central to the success of such efforts is the ability to consistently detect those features that are unique to individuals. Towards this goal, researchers are rapidly working to quantify and optimize the two primary determinants of our ability to detect individual differences - within- and between-subject variation^8–12^.

At the outset, such optimization could seem like a relatively simplistic problem, as there are only two terms to be considered. However, meaningful detection of individual differences requires optimization of the ratio of these two sources of subject-related variation (between, within), rather than the two terms independently^5,9,12,13^. Optimization of such ratios can be challenging, particularly when: i) the data necessary to address both terms (e.g. test-retest data) is either not considered or not available, and ii) there are a large number of potential contributors to each term^2,11,13,14^. A growing number of studies in neuroimaging have shifted their focus to the delineation of individual-level effects and their variations. The most immediate challenge that has become obvious is the cross-sectional nature of the vast majority of datasets, allowing for estimation of between-subject variance but lacking the repeated measurements required to establish within-subject variability. A small subset of investigators has worked to amass densely sampled test-retest datasets which can be used to assess both sources of variance^15–18^. This is not necessarily a complete solution, as one must assess the generalizability of such datasets to the larger body of data, which can be challenging due to a lack of standardized data collection protocols, as well as considerations related to sampling^19,20^. Regardless, it is a critical step towards understanding the factors contributing to the detection and optimization of individual differences, and motivating greater consideration of these issues in future study designs.

Having the data needed for optimization available, the present Perspective attempts to provide guidance for how to assess the impact of interventions on the optimization of individual difference detection. We first illustrate the impact of within-individual variation when measuring individual differences of brain-behavior associations. We then discuss sources of the within- and between-individual variation, as well as interactive effects in the design of neuroimaging experiments. Next, we review the concepts of fingerprinting, reliability, and how these two concepts can be bridged in a unifying two-dimensional space to quantify individual differences^11,21,22^. We highlight a potential analytic pitfall that can lead to contaminated estimates of inter-individual differences and enhance recently developed individual variation field maps to more accurately reflect within- and between-subject contributions. We propose the gradient flow map to guide efforts for optimizing measurement of individual differences (i.e., optimization of within vs. between-subject variation). Finally, we demonstrate the applicability of gradient flow maps for optimizing analytic strategies using fMRI-derived functional connectivity matrices.

## Within-individual Variation

### “No man ever steps in the same river twice” - Heraclitus

The human brain is a dynamic system, continuously updating itself in response to internally and externally generated stimuli and task demands in support of higher-order cognition^23^. From moment to moment, patterns of neural activity traverse the connectome, like waves moving along a river bed that is being remodeled over time. This notion is reflected in classical (psychometric) test-theory, which asserts that no two moments in time are ever exactly the same for an individual - a reality that challenges efforts seeking to meaningfully measure individual brain function^24^. Any experiment merely captures a sample of a certain time-scale moment from the entire time domain of the given individual; considered in the present context, within-individual variation is the rule — not the exception. However, in the midst of this change is a dynamic equilibrium supported by homeostatic processes. It is this equilibrium that allows the brain to maintain order despite its dynamic nature, and allows brain measures to approach repeatability over time (i.e., agreement between temporally independent test results), despite never being exactly the same^15,17,25,26^.

Research methods in experiment design and statistics have developed a variety of paradigms to control or study the within-individual variation (e.g., repeated measure, longitudinal design, etc.). However, in cross-sectional studies, within-individual variation is often overlooked or misinterpreted as inter-individual variation when studying individual differences in brain function. We demonstrate this by simulating a simple illustrative example in which subjects with known “ground truth” and inter-individual differences can generate distinct observed individual differences, driven by within-individual variation (Fig 1). In Fig 1a, we simulated a “ground-truth” score for each of the ten subjects. In an ideal scenario, we expect to obtain the ground-truth individual difference (i.e. inter-individual distance matrix, Fig 1c, left matrix). However, due to the variation of the measurement within each individual, the observed score from each test of the individual varies. As such, the resulting observed individual difference is different (Fig 1c, test 1 vs. test 2). As the variation of within-individual measurement increases (e.g., more variation across measurements, Fig 1b), the observed individual differences between tests would vary even more (Fig 1d, test 1 vs. test 2) and likely be more distinct from the “ground truth” differences. Such variation of brain measurements within individuals can also have an impact on studies of brain-wide associations. In Fig 1e-f, we show examples of varied correlation scores purely driven by the variations of measures within each sampling individual. The simulated correlation with another variable (e.g., IQ) based on the same ground-truth (r=0.35 from the population) can either result: 1) in inconsistencies in findings (Fig 1e) across samples (i.e., insignificant (r=0.282, p=0.101, n=35) or significant (r=0.365, p=0.031, n=35)) or 2) lower the statistical power to detect the brain-wise association when individual within-individual variation is sufficiently high (Fig 1f). Thus, although the ground-truths remain the same, the measured individual difference can be different due to within-individual variation alone.

**Fig 1.**
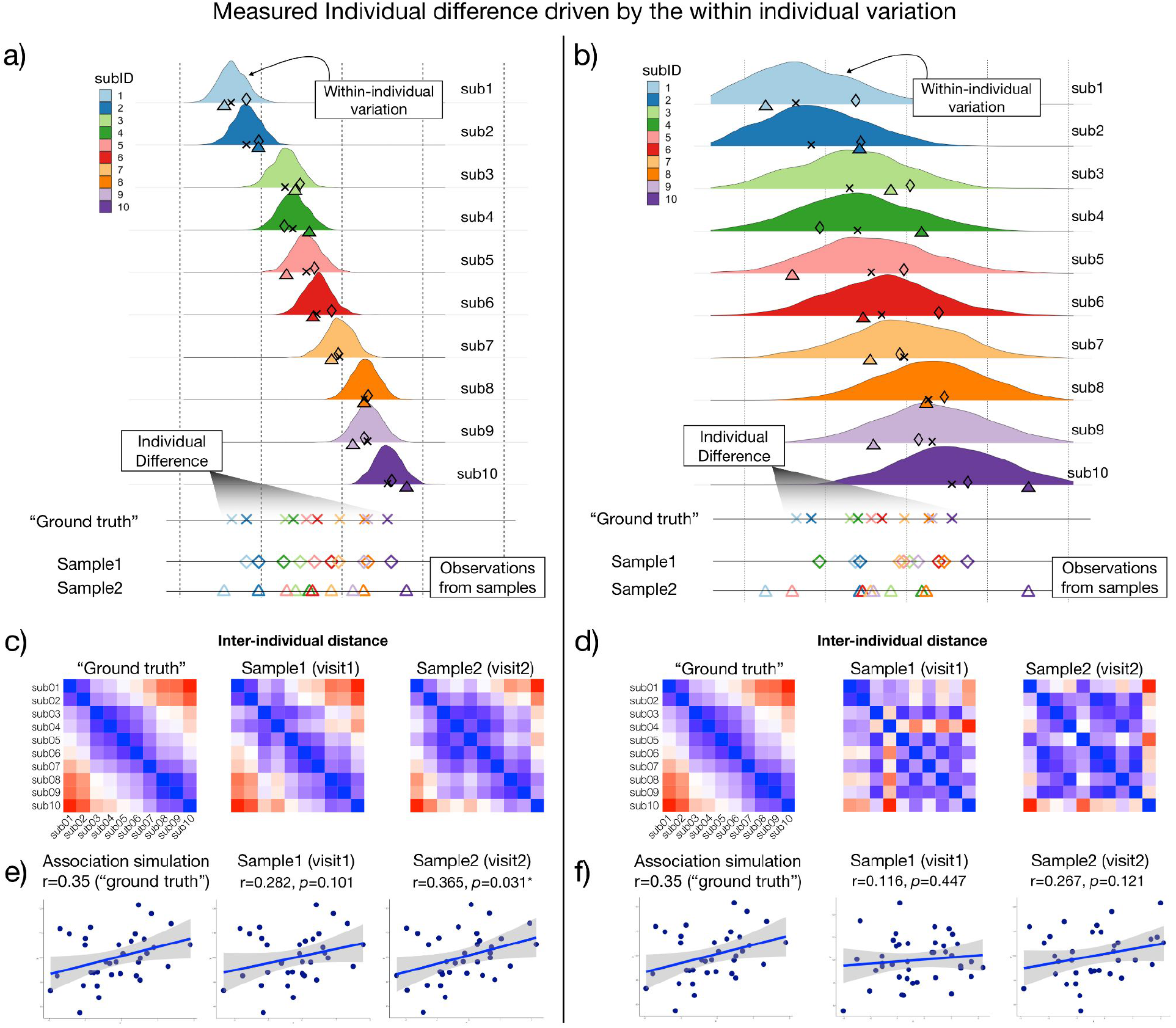
Simulations demonstrating the effects of within-individual variation on measuring individual differences. The observed individual differences can vary due to the within-individual variation. All simulations of observed inter-individual differences are generated from the same known “ground truth” (marked in cross “X”). **a–b)** Simulations of two measurements (sample1 from the first test: diamond, sample 2 from the second test: triangles) from 10 subjects with low (left panel) and high (right panel) within-individual variation. **c–d)** The measured individual differences (inter-individual distance matrix) from two measurements (sample 1 and sample 2) vary due to the within-individual variation. e–f) Associations with another simulated variable vary due to the within-individual variations. The greater within-individual variation (right panel vs. left panel) introduces noisier samples.

## Sources of Individual Variation

There are various sources of the within-individual variation when measuring the individual difference in brain function (Box 1). As depicted in Figure 2, regardless of the time-scale one considers (e.g., seconds, minutes, days, months, years), numerous potential sources of within-subject variation exist which are capable of individually or jointly impacting brain function and/or its measurements. The breadth of sources of variation can be overwhelming, though consistent with prior suggestions, they can be sorted into subgroups, such as acquisition-, experimental-, environmental-, participant-, and analysis-related factors^14,27–29^. Most of these factors are a nuisance, creating undesirable perturbations in analysis and leading to potential biases in measuring individual differences, especially when the factors are systematic in nature^30,31^. However, some can create opportunities for understanding the impact of biologically meaningful processes on brain function (e.g., time of day) when explicitly considered in the experimental design or analysis (e.g., the MyConnectome project)^15,25^.

**Fig 2.**
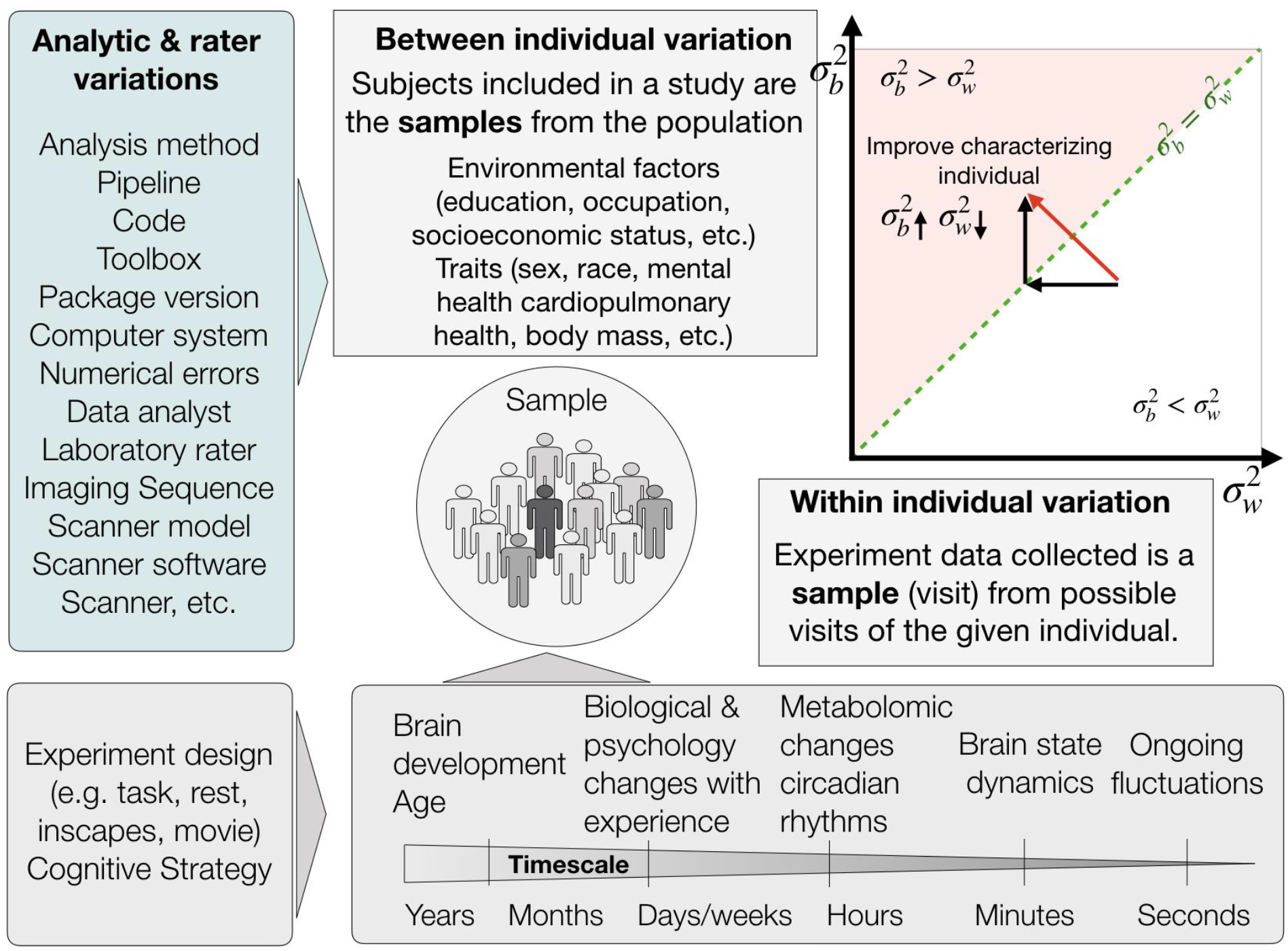
Factors affect the measure of individual differences.

How do the different sources of variation impact the measurement of individual differences? In general, measurements that have relatively lower within-subject variation and higher between-subject variation lead to improved individual differentiation^5^. It is important to note that neither the within- or between-subject variation alone determine the individual difference, but their ratio. For example, there is always variation from session to session within each individual. However, when the functional connectivity of a given subject is relatively more similar from one session to the next as compared to other individuals, this subject can be recognized and differentiated from others. When measuring individual differences of the functional connectome on a finer time-scale (e.g., functional dynamics in minutes or seconds), the dynamic changes from moment to moment may not be stable within-subject. Yet, the dynamic characteristics (e.g., the principle that governs the dynamics, how the network configurations vary, etc.) may be relatively similar within subjects as compared to across subjects^26^. Thus, it is important to decipher sources of variation at different time scales both within- and between-subjects, as well as their proportional impact in overall variations^17^.

It’s worth noting that sources of within-individual variation can also contribute to between-individual variations. For example, if participants included in the analysis are scanned at different times during the day, or undertake different experiment states (e.g., rest, task, movie)^4,32^. Even for the same task, participants can adopt different cognitive strategies resulting in variation between subjects^1^. Although previous studies have demonstrated that functional networks are largely stable within individuals, that is, variation across sessions or tasks contributes less compared to between-individual variations^17^, few studies differentiate within-individual variation from total observed variation. In particular, in cross-sectional studies, an individual’s connectome is often assumed to be stable and interpreted as part of the interindividual difference. This is problematic because as shown in Figure 1, within-individual variation can jeopardize the estimation of true inter-individual differences when it is large relative to between-individual differences. Such contaminations of between-subject differences can compromise brain-behavior association discovery.

#### Box 1 Multi-level sources of variation within individuals

Depending on the timescale of the data, sources of variation can exist across multiple levels: from moment to moment, day-to-day, month-to-month, or year-to-year. At each timescale, brain function can also vary with internal and external factors. Fig 2 summarizes common potential sources of within-individual variation in brain function. From moment to moment, brain functional organization varies along with ongoing fluctuation for each individual subject^33^. Such dynamic activity reflects the changes of the brain state with internal awareness and/or external stimulus. For example, during a resting-state scan, the brain state tends to change as the mind wanders while a subject watching a movie or undertaking a specific task (e.g., Wisconsin Card Sorting Test) will undergo state changes consistent with the ongoing stimulus^34,35^. From hour to hour, diurnal rhythms during the day can alter the brain state, as well as circadian rhythms which govern the homeostatic metabolomic and have shown daily variations on functional connectomes^36^. These can be further affected by external factors (e.g., feeding, caffeination, sleep quality the day before the scan)^15,32^. From weeks to months, seasonal effects and the life events might change the subjects’ mood and cognitive functions, affecting the brain state during measurement^37,38^. Biological and psychological changes with individual unique experiences also lead to variations in functional organization^15^. From year to year, age effects of brain development combined with long-term environmental-social influences (e.g., education, income) contribute to the variation within individuals^39–42^.

## Assessing the impact of within- and between-individual variation on measuring individual differences: from fingerprinting to reliability

To put the contributions of within- and between-individual variation to measurements of individual differences into context with one another, it is helpful to characterize the sources of variation in a two-dimensional space in which each is treated as an independent dimension (i.e. x-axis: within-individual variation, y-axis: between-individual variation)^8,9^. Within this space, conditions in which within-subject variation is low and between-subject variation is high are optimal for the detection of differences among individuals. In contrast, conditions, where between-subject variation is low and within-subject variation is high, are the least optimal. It is common practice for summarizing the relative contributions of the two dimensions in a single value. For example, when looking at continuous measurements, the ratio of between-individual variation divided by the sum of within-individual and between-individual variation, namely the intraclass correlation (ICC), is widely used to quantify how well the measure can characterize reliable individual difference^43–45^. As modern neuroscience has increased the dimensionality of characterizations for an individual, the field has faced the challenge of how to achieve such indices for multivariate profiles^22,46^. One solution that has emerged in neuroimaging is fingerprinting, a nonparametric index that quantifies whether the individuals can be matched with themselves across repetitions^21^. Alternatively, three approaches to generalizing the ICC to multivariate formulations have emerged, including i) a parametric extension of the classic ICC formulation, image intraclass correlation coefficient (I2C2)^47^, ii) distance-based ICC (dbICC), which reformulates the ICC in terms of distances^48^, and iii) discriminability, a nonparametric index that assesses the degree to which an individual’s repetitions are relatively similar to one another^22,49^.

#### Box 2 Indexing individual difference

The toy example includes two repetitions in ten participants. Figure 3a shows the pairwise symmetric distance matrix between each of the repetitions, ordered by participants. The block-diagonal elements indicate how different the repeated measures are within participants, whereas off-diagonal elements represent the dissimilarity of measures between participants. We then construct a two-dimensional space by plotting the observed between-individual elements along the y-axis against the within-individual value on the x-axis (Fig 3b). Each between-individual element for a row or column in the distance matrix can be represented by an array of dots in the distance space. It’s important to note that the x=y line in this space represents the case that within-individual distance equals the observed between-individual distance. The dots above the x=y line (i.e. x<y, red shaded zone) represent the cases that the difference within individuals is lower than the difference observed between individuals. By counting the number of dot-columns that fully fall above the x<y zone (i.e. within-individual distance < between-individual distance) as compared to the number of individuals (i.e. total number of dot columns), we can calculate the percentage that the individual subjects can be successfully differentiated from the group (i.e. identification rate, a.k.a. fingerprinting). In the toy example, 3 out of 10 dot columns are above the x=y line and resulting in a fingerprinting score of 0.3 (Fig 3b). If more dots fall above the x<y zone, the repetitions within individuals are more likely similar to one another but distant from other individuals. Instead of counting the percentage of the dot columns above the x=y line, discriminability measures the percentage of dots above the x=y line as compared to the total number of dots. We can also estimate the ratio of between-individual variation in total observed individual variation (i.e. ICC, equations in Fig 3b).

In neuroimaging and psychology, ICC and its extension (e.g. I2C2, dbICC) are widely used approaches for assessing reliability (i.e. the degree of agreement or consistency of measures)^47,48,50,51^. In the context of measuring individual differences, ICC, by definition, represents how much inter-individual variation can be measured. This is particularly useful in studying individual differences and brain-behavior relationships. For example, if ICC=0.5 of a given measurement, regardless of neuroimaging or behavioral measurement, indicates that the observed measurement can only capture 50% of the inter-individual variation of interest from observed inter-individual variation that is contaminated with within-individual variations.

**Fig 3.**
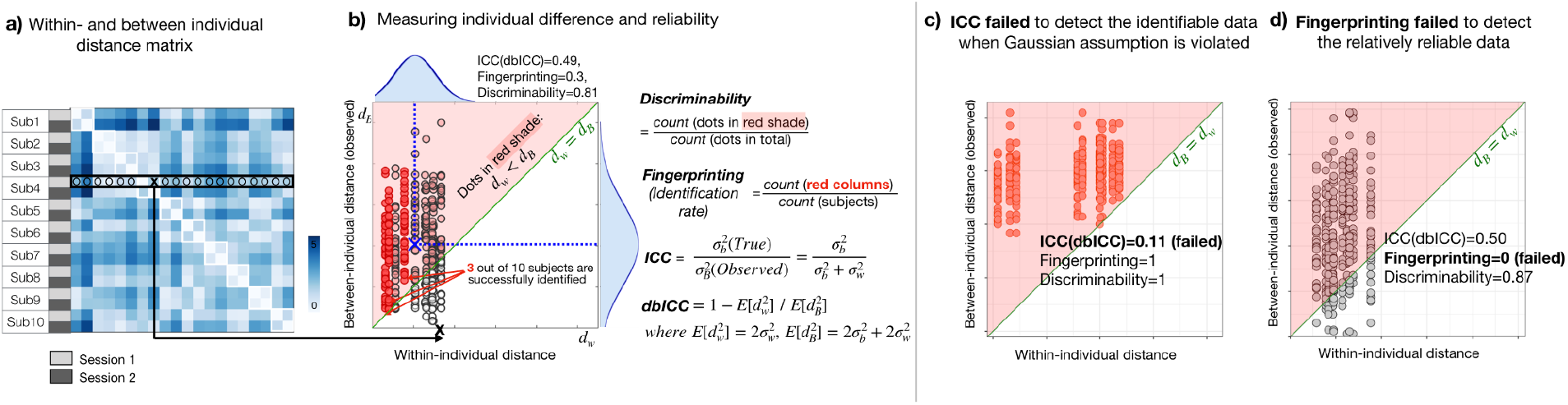
The two-dimensional distance field map characterizes within- and between-individual variability. **a)** Distance matrix across individuals and two repeated measures. Diagonal submatrices depict within-individual distances, while off-diagonal elements are between-individual distances **b)** Identifying individuals are quantified by discriminability, intraclass correlation (ICC), or fingerprinting (i.e. identification rate). Discriminability is estimated by the probability that the distance between sessions within each individual is smaller than distances to other individuals. ICC values are estimated with the ratio of between-individual variation divided by the sum of within-individual and between-individual variation. Fingerprinting is estimated by the proportion of individuals correctly identified (modified from^11^). **c-d)** Examples in which fingerprinting and ICC are essentially uninformative and arguably provide misleading information.

With many indices emerging to take on the challenge of measuring individual differences and calculating reliability, understanding the advantages and limitations of each can aid in selecting a few in applications. First, we draw attention to parametric assumptions for a given data set (e.g., Gaussian distribution, homogeneous variance). In cases where data is not Gaussian distributed, ICC and dbICC can be highly misleading (Fig 3c). Additionally, in some cases, which often occur in neuroimaging studies, ICC and dbICC are negative due to the negative difference between two mean-square terms in the computational formula^43^; although not inherently problematic, in practice, negative ICC/dbICC is not interpretable. Second, each index provides different sensitivity. High discriminability is required for fingerprinting (i.e. identification), but not vice versa; in some conditions, fingerprinting and discriminability will diverge, with fingerprinting potentially leading to the wrong conclusions^11^. To understand this, we draw your attention to a simple scenario using a toy model in Fig 3d. Specifically, situations can arise where most, but not all, of the between-individual distances are larger than the within-individual distance (i.e. most dots are in the x < y zone, but none of the entire dot columns are in x < y zone; x = within-subject distance; y = between-subject distance). In such instances (Fig 3d), the individual difference is relatively discriminable, but fingerprinting would be zero — and thus, misleading with respect to potential for optimization and eventual usage.

An important caveat to be aware of when using multivariate indices of reliability is that they do not guarantee the reliability of each univariate feature. It is well established that reliability differs across regions and connections in the brain^50,52^. Some features may contribute more to the detectability of individual differences than others^53^; and, it is possible that some features may differentiate a subset of individuals, but not all. As such, the reliability for a multivariate profile should not be assumed to be a reflection of that for its individual features. A sensitivity test (e.g., leave-one-out analysis) to examine the contributions for each of the individual features or the univariate reliability needs to be considered.

## Optimizing Measurement of Individual Variation

Once one has properly assessed reliability for a given measure, the next logical question is: how can reliability be improved. While researchers commonly focus on reliability alone in optimization efforts, this can be problematic due to its ratio nature of two individual-related sources (within and between-subject variation). As with any ratio measure, the best intuitions for efficient optimization require one to take into account each of the terms individually as well as in combination. Here, we formalize a framework for assisting researchers in this process, referred to as “gradient flow mapping” and introduce a supporting online tool - Reliability Explorer (https://tingsterx.shinyapps.io/ReliabilityExplorer/). Central to the concept of gradient flow mapping is to construct the variation space (Fig 4a-c) using the estimated “true” between-individual variation 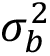 (y-axis) against the within-individual variation 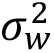 (x-axis). Unlike the distance space above (Fig 3), here we focus on the estimated “true” between-individual variation rather than the observed between-individual distance. This distinction is important because, as previously discussed, the observed between-subject variation is contaminated by the within, and resulting individual differences are overestimated. Separating “true” between-individual variation and within-individual variation enables understanding the contribution of each term for optimization.

**Fig 4.**
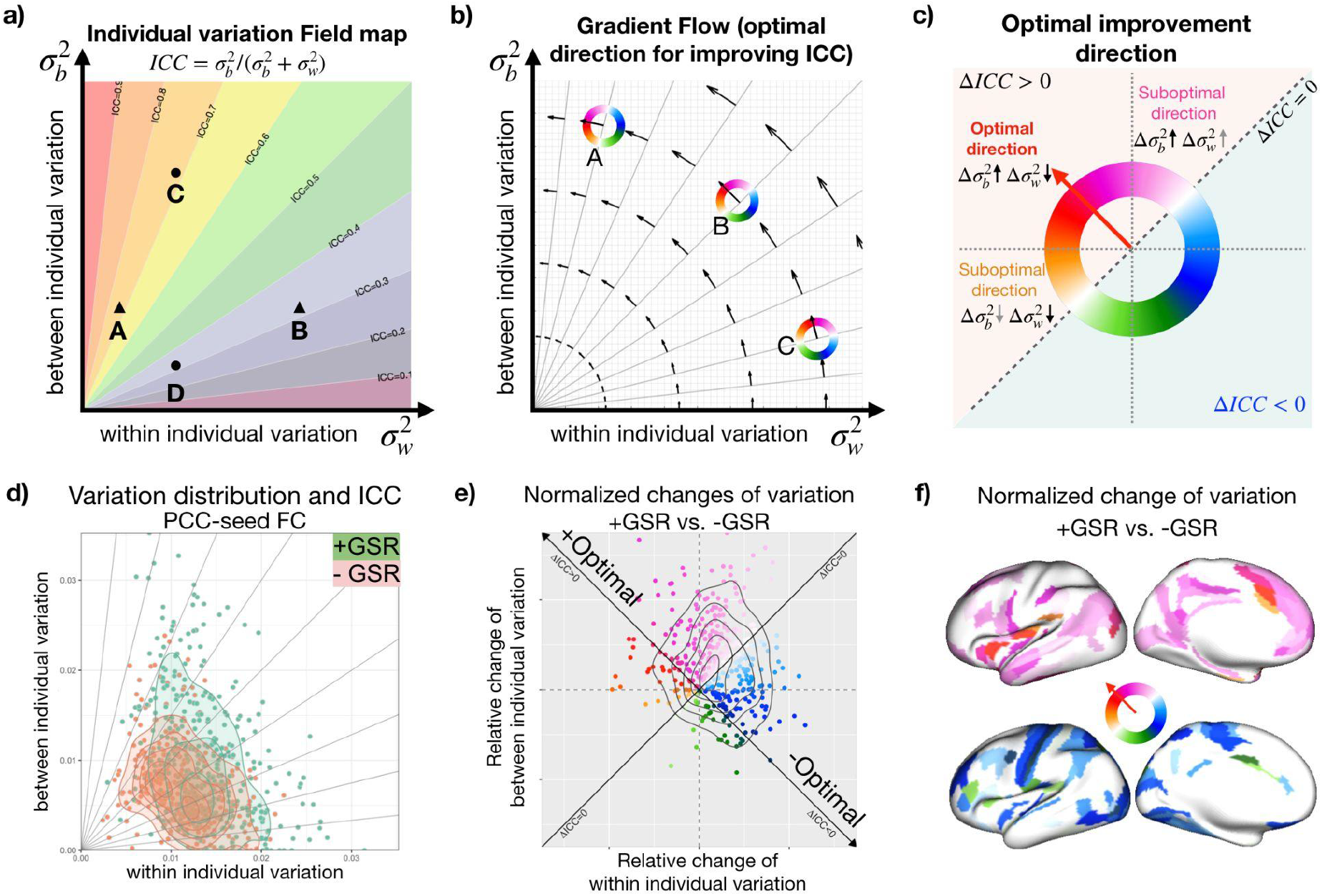
Individual variation field and its gradient flow map guide the comparison and optimization of measuring individual differences. **a**) The two-dimension theoretical individual variation field map characterizes within- and between-individual variability and the likelihood of individual characterization (quantified by the intraclass correlation reliability). **b)** The theoretical gradient flow map captures the most efficient way for improving reliability (i.e. the first derivative of the ICC). **c)** Normalized changes of variation as compared to the optimal direction for improving ICC. **d)** The within- and between-individual variability of functional connectivity (seed: posterior cingulate cortex [PCC]) calculated based on two preprocessing pipelines (with and without GSR). **e**) Relative change of variation normalized along to the optimal direction. **f**) Relative change of variation mapped on the cortex.

As shown in Figure 4a, a measurement with lower within-individual variation (i.e. decrease x, from point B to point A) and higher between-individual variation (i.e. increase y, from point D to point C) provides higher reliability (i.e. ICC). However, contributions of the within- and between-individual variation for improving the reliability are not always the same. For example, if a measure has relatively small x and large y (e.g. point A in Fig 4b), the reduction in x (within-individual variation) improves ICC more than a similar increment in y (between-individual variation). On the other hand, if a measure is relatively high in x but low in y (e.g. point C in Fig 4b), the most efficient direction to improve the reliability is to increase the between-individual variation. Such optimal direction for improving reliability can be calculated as the first derivative of the ICC (Fig 4b). The improvement of x and y that is closest to this optimal direction is more likely to improve the discriminability the most under Gaussian assumption. Additionally, by normalizing the changes of the ICC as compared to the optimal direction (Fig 4c), we can assess how possible analytic strategies and experiment designs improve the reliability and whether the improvement is in the most efficient direction. In Figure 4d-e, we showed an example of comparing the reliability of functional connectivity (seed: posterior cingulate cortex) between two analytic pipelines (with and without global signal regression [GSR]). Utilizing the variation field space and the gradient flow map, it’s easier to understand how different analytic methods manipulate the within- and between-individual variation and whether it improves the reliability to characterize individual differences.

## Concluding Remarks

Reliability of measurement is a well-established prerequisite of individual difference research. However, when faced with the challenges of generating the necessary datasets to meaningfully estimate individual differences, many psychology and neuroscience studies have forged ahead without careful consideration of within-individual variation or its relevance to measurement reliability. This work provides a critical guide for conceptualizing reliability in terms of its component variations (within- and between-individual variations). We drew attention to the reality that within-individual variation, regardless of whether it is meaningful or noise, is always embedded in the observed scores of individual differences. We highlighted measurement reliability as a critical means of accounting for such variations and putting them into context with between-individual variations, providing insights into how to best choose a method for quantifying reliability for a given analytic context. Additionally, we updated the calculation approach for reliability field maps to better account for within-subject contributions to individual differences.

Recognizing the growing need for techniques to guide optimization efforts for measurement of individual differences, we further proposed the reliability gradient flow map to quantify the optimization efforts of measuring reliability and individual variations. In support of our Perspective, we develop an online app that integrates reliability concepts, calculation, and optimization to bridge the gap between establishing reliability and measuring individual variations (https://tingsterx.shinyapps.io/ReliabilityExplorer). We hope that the guidance and toolbox will help calculate and compare reliabilities across experiments and analytic methods to facilitate studying individual differences in neuroscience and psychology.

## Code Availability

The toolbox is implemented using web-based R/Shiny application and is freely available on shinyapps.io: https://tingsterx.shinyapps.io/ReliabilityExplorer

## Acknowledgments

This work is supported by gifts from Joseph P. Healey, Phyllis Green, and Randolph Cowen to the Child Mind Institute and the National Institutes of Health fundings (R24MH114806 to MPM, RF1MH128696 to TX, Additional grant support for JTV comes from R01MH120482 (to Theodore D. Satterthwaite, MPM) and funding from Microsoft Research.

